# Deciphering 3D Human Sphygmopalpation Pulse Patterns using “X-ray” Images Acquired from Tactile Robotic Fingers

**DOI:** 10.1101/2021.01.10.426152

**Authors:** Ka Wai Kong, Ho-yin Chan, Jun Xie, Francis Chee Shuen Lee, Alice Yeuk Lan Leung, Binghe Guan, Jiangang Shen, Vivian Chi-Woon Taam Wong, Wen J. Li

**Affiliations:** City University of Hong Kong, Department of Mechanical Engineering, Hong Kong; The University of Hong Kong, School of Chinese Medicine, Hong Kong; Advanced Inkjet Systems Co., Limited, Taiwan

## Abstract

*Sphygmopalpation* at specific locations of human wrists has been used as a medical measurement technique in China since the Han Dynasty (202 BC - 220 AD); it is now generally accepted that traditional Chinese medicine (TCM) doctors are able to decipher 28 types of basic pulse patterns using their fingertips. This TCM technique of examining individual arterial pulses by palpation has undergone an upsurge recently in popularity as a low-cost and non-invasive diagnostic technique for monitoring patient health status. We have developed a pulse sensing platform for studying and digitalizing arterial pulse patterns via a TCM approach. This platform consists of a robotic hand with three fingers for pulse measurement and an artificial neural network (ANN) together with pulse signal preprocessing for pulse pattern recognition. The platforms previously reported by other research groups or marketed commercially exhibit one or more of the following imperfections: a single channel for data acquisition, low sensitivity and rigid sensors, lack of control of the applied pressure, and in many reported works, lack of an intelligent data analysis system. The platform presented here features up to three-dimensional (3D) tactile sensing channels for recording data and uses highly sensitive capacitive MEMS (microelectromechanical systems) flexible sensing arrays, pressure-feedback-controlled robotic fingers, and machine learning algorithms. We also proposed a methodology of obtaining “X-ray” image of pulse information constructed based on the sensing data from 3 locations and 3 applied pressures (i.e., mimicking TCM doctors), which contains all arterial pulse information in both spatial and temporal spans, and which could be used as an input to a deep learning algorithm. By applying our developed platform and algorithms, 3 types of consistent pulse patterns, i.e., “Hua” 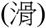, “Xi” 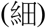, and “Chen” 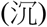, as described by TCM doctors”, could be identified in a selected group of 3 subjects who were diagnosed by TCM practitioners. We have shown the classification rates is 98.7% in training process and 84.2% in testing result for these 3 basic pulse patterns. The high classification rate of the developed platform could lead to further development of a high-level artificial intelligence system incorporating knowledge from TCM – the robotics finger system could become a standard clinical equipment for digitalizing and visualizing human arterial pulses

## Introduction

Traditional Chinese Medicine (TCM) has been used for healthcare in China for more than two thousand years. TCM physicians use four diagnostic methods including inspection, auscultation and olfaction, inquiry and palpation to collect clinical information in order to make diagnosis for the constitution and syndrome pattern recognition. TCM sphygmopalpation (TCMS) [1], a combination of human arterial pulse sensing and diagnosis, has been used by TCM physicians since the Han Dynasty (202 BC - 220 AD). Different than Western medicine practitioners, which use the palpation for estimating cardiovascular functions based on the pulse rates and rhythm, experienced Chinese medicine practitioners (CMPs) can use their fingers’ sensations and their own experience to draw conclusions about patients’ holistic health status. According to the theory of TCM [1] and classical TCM concepts recorded in an ancient Chinese masterpiece called “Mai Jing” (“The Pulse Classic”) [2], arterial pulses detected at three different locations (i.e., CUN, GUAN and CHI) of both wrists reflect the health conditions of the internal organs, “Qi-Blood” and “Yin-Yang status”. Qi-Blood is an inseparable term in TCM. Qi has been translated as “vital energy”, which is generated by the internal organs, to nourish human body through the movement of Blood, which is a denser form of and moved by Qi. TCM describes Yin-Yang as relative opposite qualities or manifestations of Qi. If there is any change of physiological states in the internal organs and the related functions, the status of the pulse will be affected, forming its unique diagnostic basis. Experienced TCM physicians have developed advanced skills to sense the changes of the pulse patterns for their diagnosis. Although there are many written rules and well-proven records of the success of TCMS, the communication of the corresponding knowledge and skill is still based on individual understanding and experience, which needs a scientific verification. In contrast to electrocardiography (ECG), which already has a standard data acquisition procedure [3], the development of TCMS standardization and arterial pulse digitization is important for reliable and consistent diagnosis. Hence, it is worthwhile and extremely critical to study TCM arterial pulses by a scientific, quantifiable and reliable approach. It is also important to collect correct pulse signals based on TCM theories and methodologies. Hence, an ideal measurement system for mimicking the fingers of CMPs must have the ability to provide different levels of pressure, measure arterial pulses precisely at the CUN, GUAN, CHI points, and classify 28 arterial pulse characteristics according to TCM theory [4], as shown in Table I.

**Table I:**
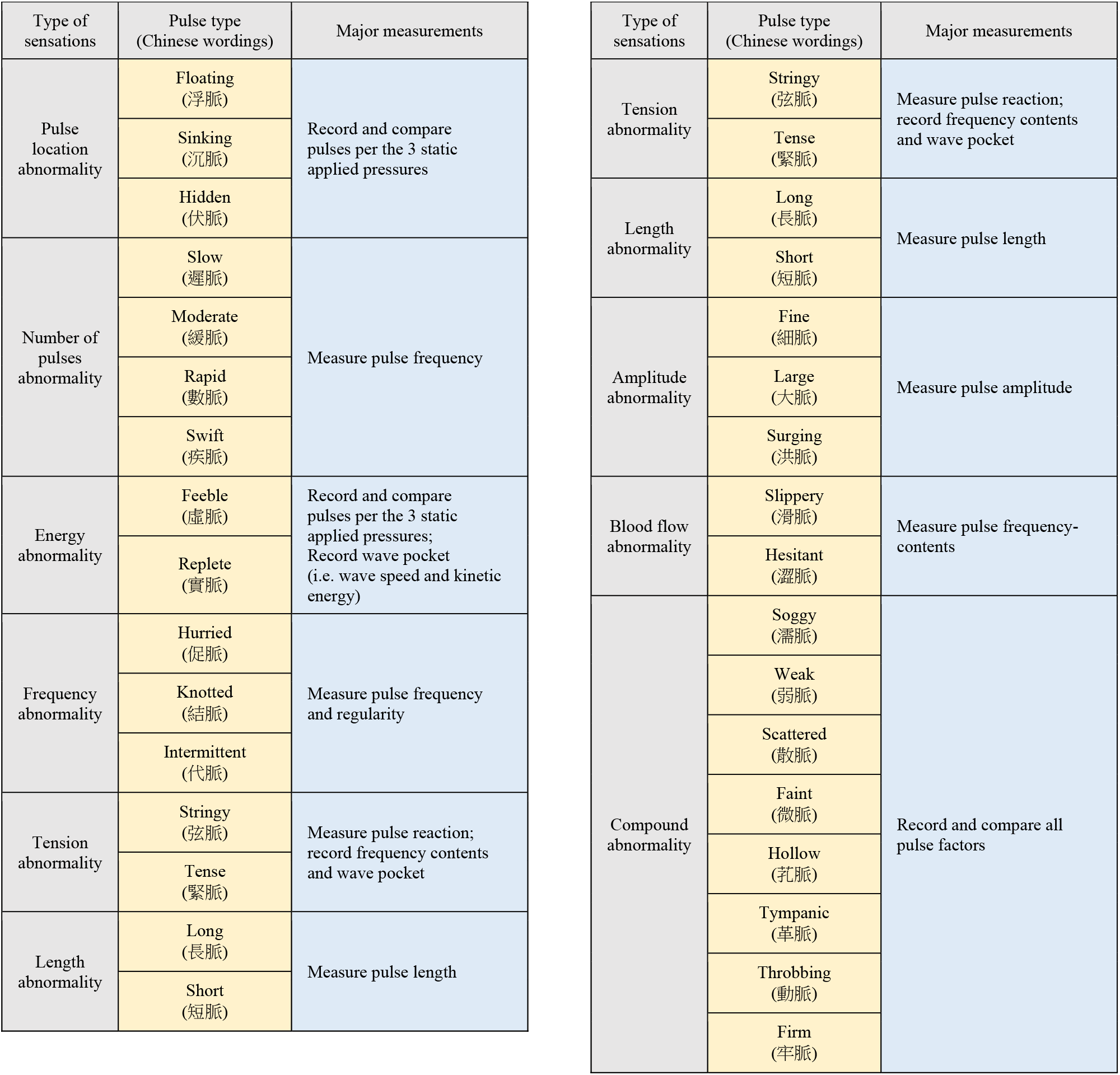
List of all recorded pulse types.

For the past twenty to thirty years, vigorous academic TCM research efforts in China [5], Taiwan [6] - [8] and Hong Kong [9] [10], as well as other locations [11], have pursued the development of useful instrumentations to capture the arterial pulse at the human wrist. These machines have often exhibited design issues that hinder their development. Some of them, for example [12], simply used a single pressure sensor for arterial pulse recording and diagnosis without considering the position where the pulse should be taken. Other machines [13]-[16] recorded pulses from different positions but still omitted the importance of the pressure applied on the wrists. However, in TCMS, a conclusive pulse diagnosis can be achieved only by analyzing all pulses measured from the three key positions (CUN/GUAN/CHI) and their variations under different applied pressures.

In addition to the development of instruments for pulse measurement, pulse pattern classification and analysis are also important for standardizing TCMS. The most common and effective technique is the use of artificial neural networks (ANNs), a machine learning method. Four important design parameters influence the performance of an ANN: data source, the size of the training sets, input features and output targets, and network structure [17] [18]. Some ANNs have been developed for TCM [19]-[22]. The source of the data collection was either a simple pulse monitoring system [23] [24] or arterial pulse reference books [25] [26]. The performance of these algorithms is restricted due to the limited amount and range of data obtained from a single patient. Additionally, none of these networks has addressed the influence of the arterial pulse under different applied pressures. These networks were developed without strictly following the written rules of TCMS; therefore, their performance may be far from that expected of TCMS.

In this paper, we discuss the development of a novel pulse sensing platform (PSP) that can record and classify arterial human pulses via the TCMS approach, which has never been achieved before. This platform can be divided into two major parts: (1) a palpation robotic hand (PRH), which consists of three robotic fingers for pulse measurement, and (2) a dedicated control and signal processing algorithm for pulse data filtering and classification. The developed system can adopt and learn from different TCM practitioners who may belong to different martial arts disciplines and have different interpretations of arterial pulses. For instance, the disciplines can be divided into Simultaneously Palpation (SP) and Pressing with One Finger (PWOF). The entire trend of body state is verified by SP, and unique characteristics of viscera and bowels are verified by PWOF [27]. With further development and big data analysis, this system can provide a conclusive pulse diagnosis according to what it has learned. This system will benefit the development of more reliable and accessible TCM by providing quantifiable sphygmopalpation arterial pulse information.

## Methods

### Design of a Palpation Robotic Hand (PRH)

The overall design of the PRH is shown in Figure 1(a). The machine consists of three robotic fingers, as shown in Figure 1(b), that are driven by three individual driving-torque motors via metal strings. The mechanical design and driving mechanism of the fingers are shown in Figure 2(a). The movement and the applied force (*F_r_*) of the finger, as shown in Figure 2(b), are achieved by the force balance between a metal string (*T_s_*) and a restoring spring (*F_k_*). A more detailed discussion can be found in this reference [27]. Each fingertip is curved similarly to a human finger and mounted with a flexible capacitive sensor, as shown in Figure 2(c). The sensors are custom-made sensors from Pressure Profile System, US [28]. Each sensor has 4 x 6 sensing elements with an element size of 2 mm x 2 mm. The working range of each element is from 0 to 9 psi with a repeatability of 0.7%. The resultant pressure can be calculated by averaging either all 24 element readings or a selection of individual sensing elements. An example of a pulse signal recorded for 5-second interval is shown in Figure 3. One of the major advantages is that an array of sensors will give better position tolerance, which reduces the positional accuracy requirements of the robotic fingers. Moreover, the sensor array can give both temporal and spatial information on a pulse. To show the actual temporal arterial pulse picked up at 3 locations (i.e., CUN, GUAN and CHI, called CGC in this paper) under 3 applied pressures (i.e., FU, ZHONG and CHEN, called FZC in this paper), the selected data were plotted using 3D color contour maps and the data selection procedure are shown in Figure 3.

**Figure 1:**
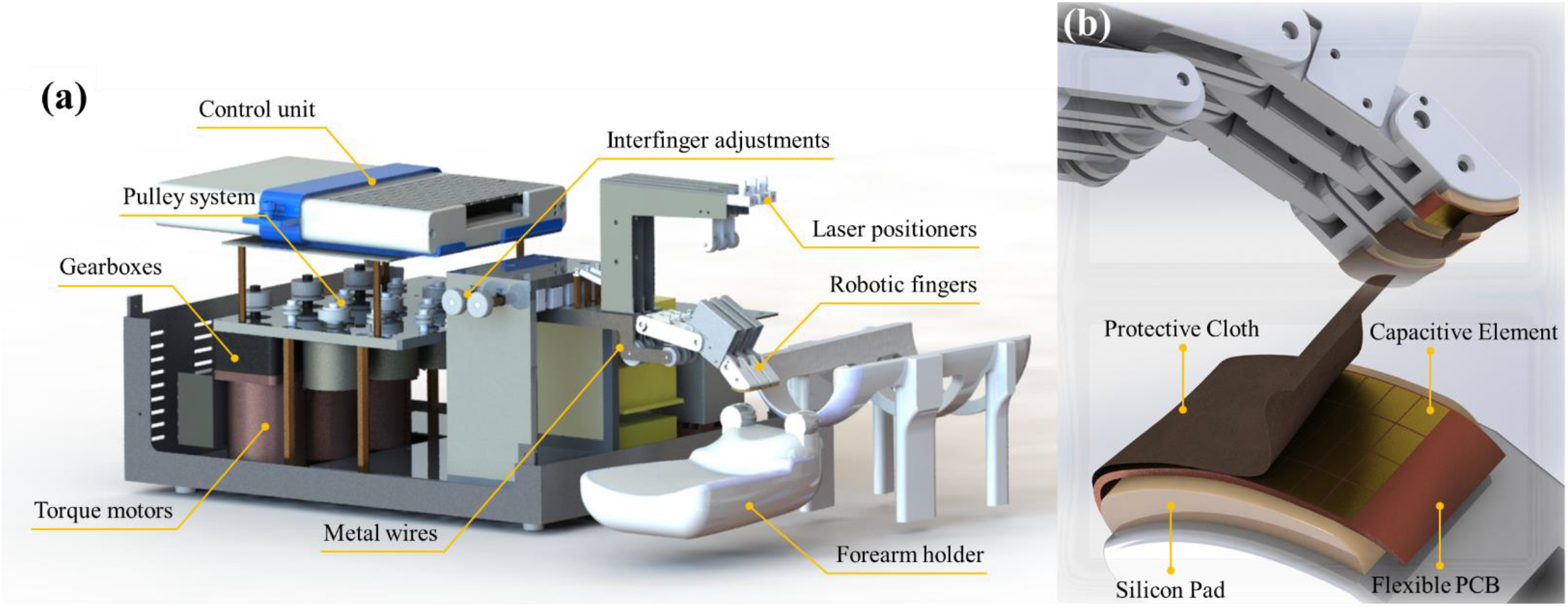
The design and setup of the pulse sensing machine: (a) The overview of the machine; (b) zoom-in of the robotic fingers and mounted tactile sensor.

**Figure 2:**
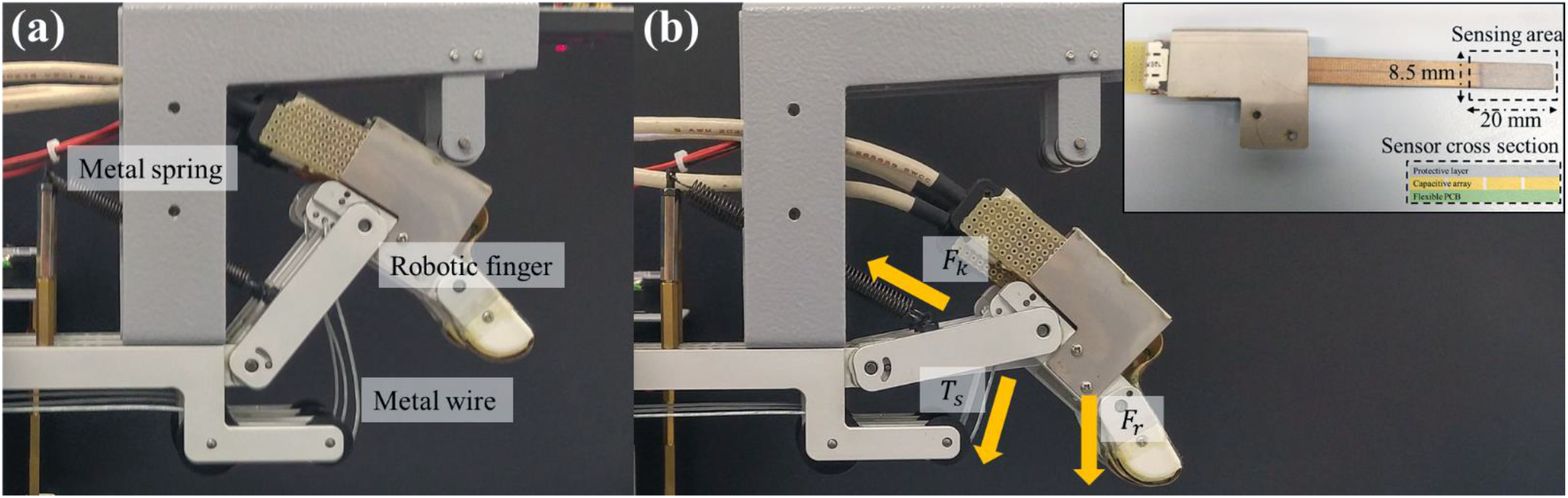
The driving mechanism of the finger: (a) Original position of the finger (F_k_=T_s_, F_r_=0); (b) the finger is fully extended (F_k_<T_s_, F_r_>0) [inset: The fingertip sensor and cross-section view of the sensing area].

**Figure 3:**
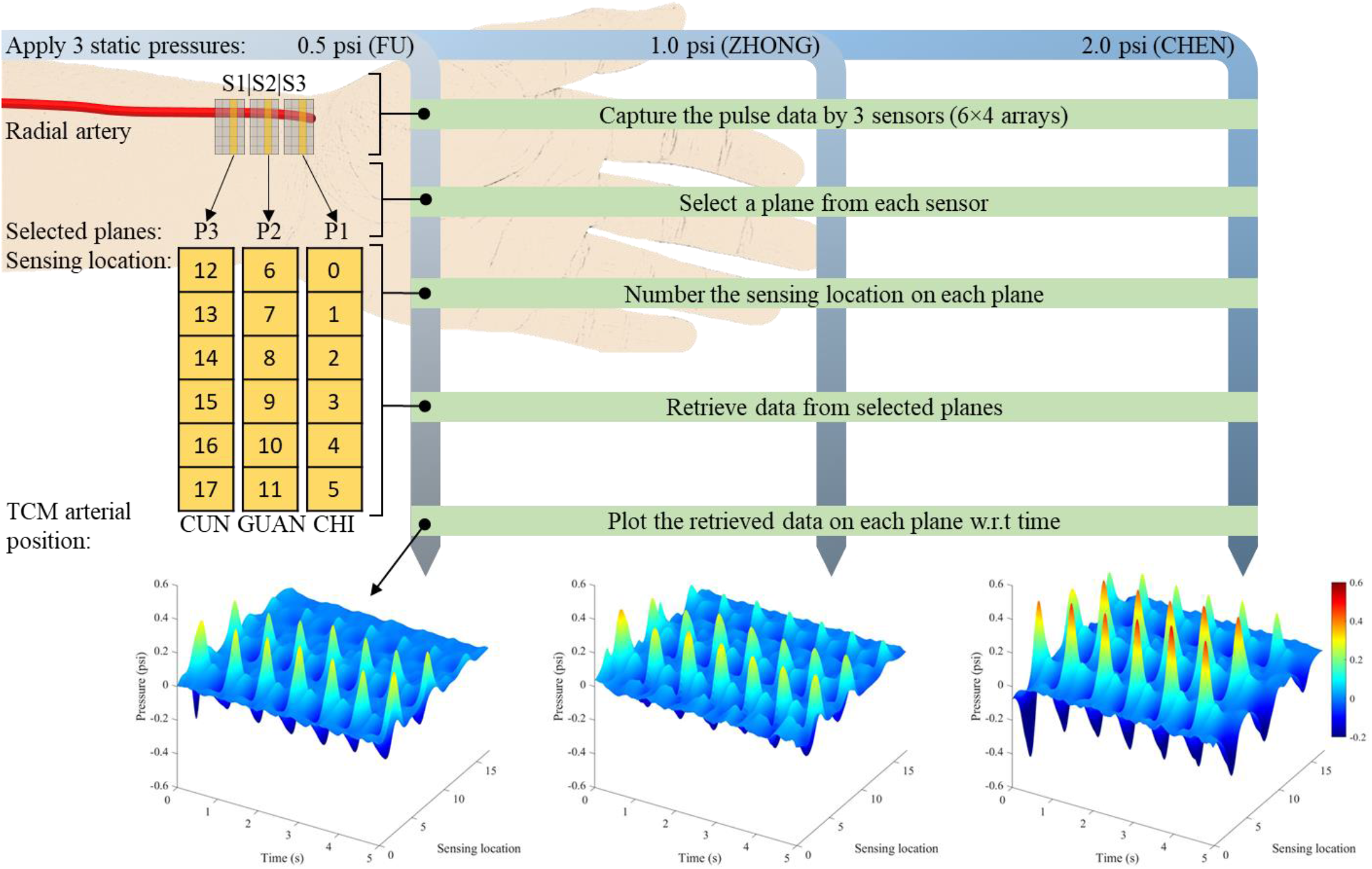
3D arterial pulse record of human wrist at the location of CUN, GUAN and CHI under 3 levels of static pressure (FU, ZHONG and CHEN).

In addition, three unique features of the machine design ensure the accuracy and repeatability of pulse measurement positions (i.e., CUN, GUAN and CHI, called CGC in this paper) on the subject’s wrist, as shown in Figure 3. First, interfinger distances can be manually adjusted. This ability is an extremely important feature since CGC positions vary among individuals. For example, a person with a shorter forearm will have CGC positions closer together. Second, three LED lasers are used to show the expected finger positions on the subject’s wrist. The task, then, is to align these 3 laser spots with the CGC positions on the wrist. Third, each finger can be individually actuated and can maintain a precise applied pressure depending on the surface topography of the skin and the various biological tissues and bones underneath.

### Control and Signal Processing Algorithm of the PSP

During palpation, CMPs usually adjust their fingertip pressures to collect additional arterial pulse information. They apply three levels of fingertip pressures, namely, FU, ZHONG and CHEN, called FZC in this paper. In this paper, we calibrated these 3 pressures based on CMP recommendations and set them to FU=0.5 psi (25.9 mmHg), ZHONG=1.0 psi (51.7 mmHg) and CHEN=2.0 psi (103.4 mmHg). Arterial pulses were recorded in at least 1-minute intervals. The control logic and signal processing flow of the PSP are shown in Figure 4. The PSP is designed to follow the technique described by a skilled CMP. The fingers are actuated by applying voltages to the torque motors. The fingertip sensor is used as an input into the feedback control loop for real-time monitoring of the fingertip pressure. Once the designated pressure is reached, the sensors start recording pulse signals from the CGC positions. Additionally, pulses will be taken at the FZC pressures. Therefore, a total of 9 combinations of arterial pulses per hand will be obtained for every subject. Together with a CMP’s diagnosis, these pulses will be input into the signal processing unit for further waveform preprocessing and classification.

**Figure 4:**
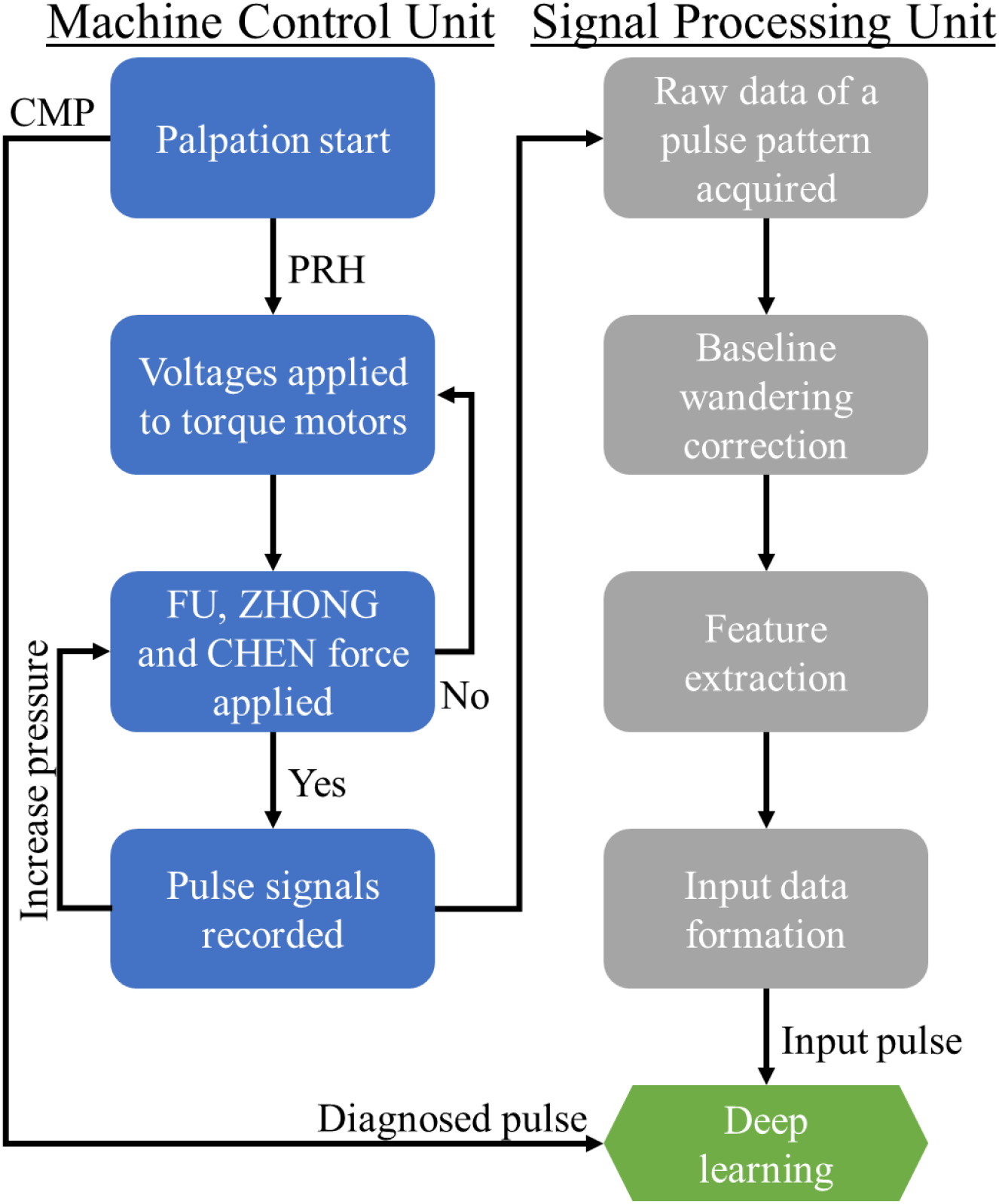
Flowchart of control logic and signal processing of the PSP.

In the signal processing unit, two signal preprocessing steps are required to generate a final input set for an ANN: baseline wandering correction, which removes undesired noise, and feature extraction, which defines a small number of feature points representing a single arterial pulse. Then, the processed data will be input into the ANN, which will be discussed later, for machine learning and pulse classification.

### Definition of an Arterial Pulse in Our System

Electrocardiography (ECG) is a common tool for diagnosing heart diseases in Western medicine. Numerous studies on ECG show convincing cardiac disease classification results obtained by using the QSR complex as feature points [29] – [32] and ECG signal classification with ANN [33] – [35]. In Figure 5(a), a normal ECG sinus rhythm is marked by the QRS complex [36], which is a combination of three deflections on a typical ECG. In contrast, the arterial pulse has two peaks (i.e., systolic and diastolic) and one notch (i.e., the dicrotic notch) on the sphygmogram, as shown in Figure 5(b). According to TCM theories, arterial pulses, which are generated by the heart pumping blood, reflect the holistic health status of human subjects. There are 28 basic well-known pulse characteristics [37], which reflect different status of holistic healthy and pathological statuses according to TCM knowledge.

**Figure 5:**
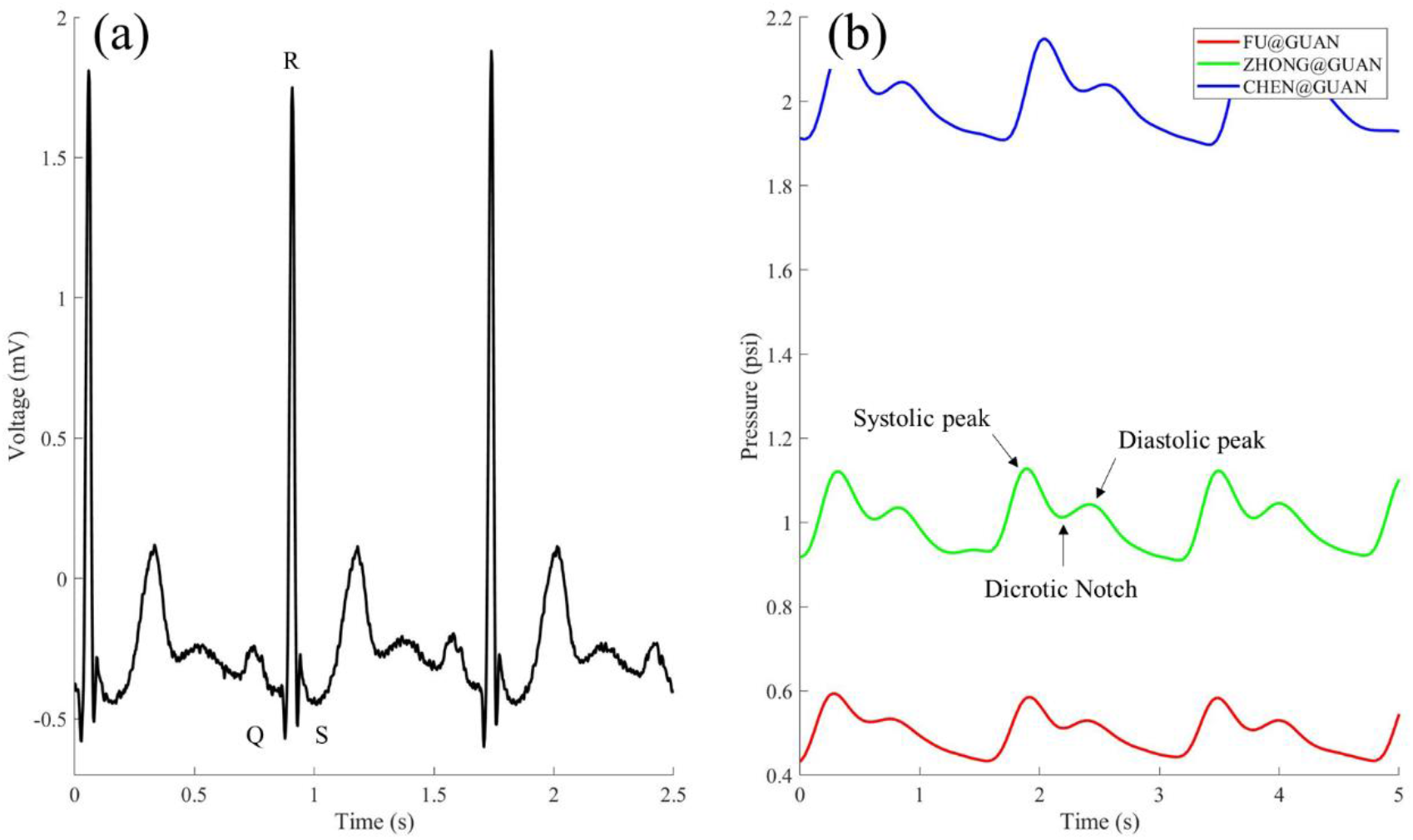
(a) A normal sinus rhythm of ECG signal from modified lead II [36]; (b) a typical arterial pulse measured from wrist under 1.0 psi (51.7 mmHg) fingertip pressure.

A normal arterial pulse means the person has normal Qi-Blood and Yin-Yang Status in TCM concept which normal cardiovascular functions and healthy condition, when the characteristics of their pulse are palpated by a CMP. In our system, due to the large amount of raw data collected, we need an effective and efficient way to reduce the data size. Therefore, each pulse was defined by five points, namely, the 2 peaks (systolic and diastolic), 1 notch (dicrotic), and the start and end points, as shown in Figure 5(b). An example of the five points is listed in Table II. The average resultant pressure at a particular time instance is calculated by taking the pressure average among all 24 sensing elements and subtracting the applied pressure *P* fingertip, as shown in the equation below:

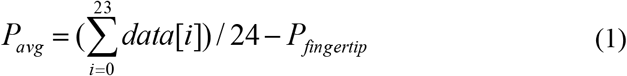

where *P_avg_* is the average pulse pressure and *data[i]* is the pressure reading of the *i*-th sensing element over a robotic finger.

**Table II:** An example of the five feature points that define a pulse in our system.

| Time (s) | Avg. pressure (psi) | Feature points |
| --- | --- | --- |
| 0.00 | -0.011 | 1<sup>st</sup> min (Start point) |
| 0.16 | 0.152 | 1<sup>st</sup> max (Systolic peak) |
| 0.30 | 0.057 | 2<sup>nd</sup> min (Dicrotic notch) |
| 0.38 | 0.078 | 2<sup>nd</sup> max (Diastolic peak) |
| 0.82 | -0.019 | End point |

### Signal Preprocessing

As shown in Figure 6(b), the measured arterial pulses have baseline drift problems. There are two main reasons for the baseline fluctuation during measurement: natural movement of the volunteer’s hand and body due to breathing and natural variation in the systolic and diastolic pressure. This baseline drift makes pattern recognition more difficult. Therefore, either the training matrix must include training vectors over a wide spread of baseline pressures, or the raw data must be processed to provide a standard baseline to the neural network. In our proposed method, the preprocessing of raw data was chosen to reduce the complexity of the input layer of the network for the classification of pulse patterns. Hence, the raw data were filtered and rearranged.

**Figure 6:**
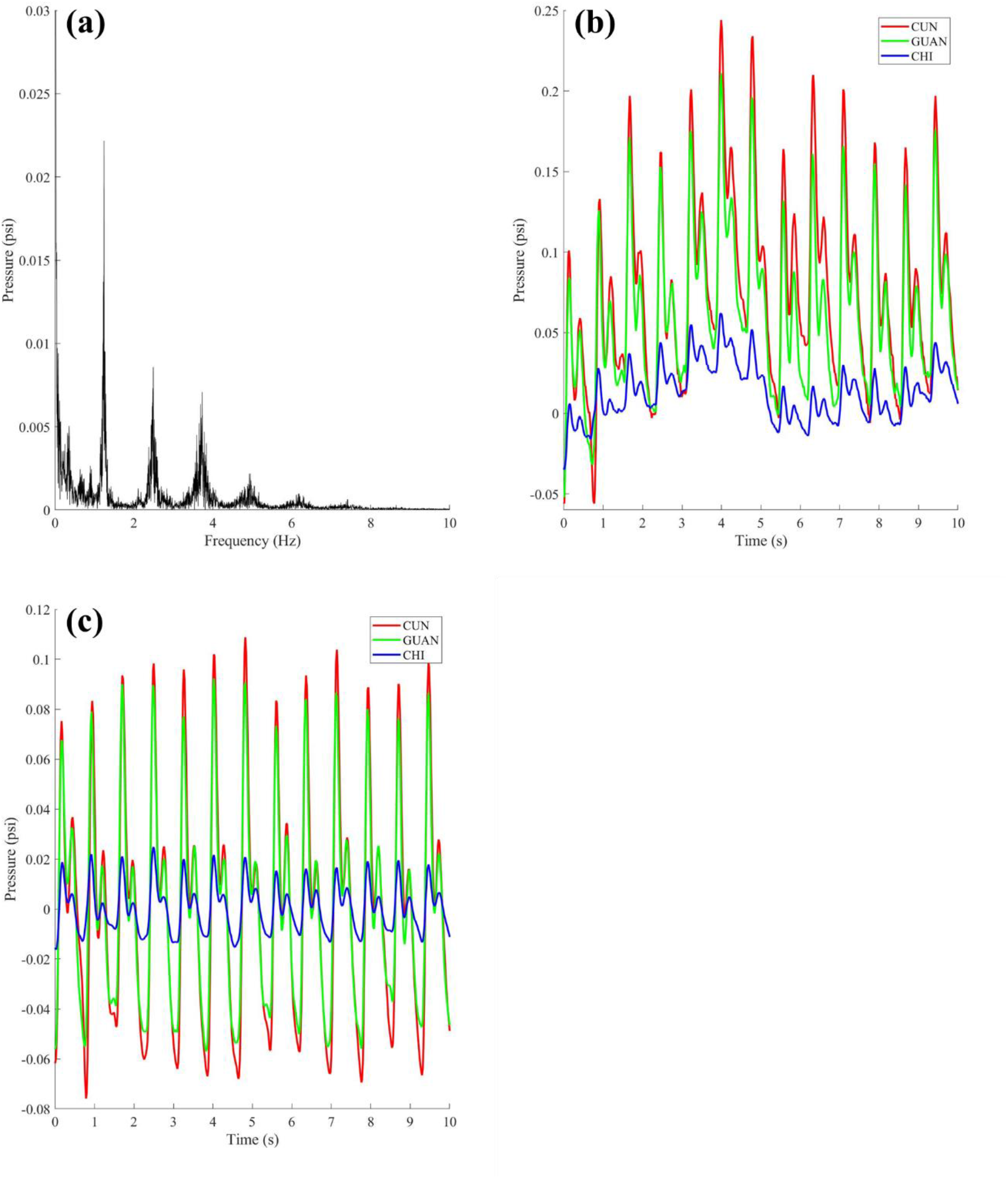
Signal processing for pulse data of subject 1. (a) An example of Fast Fourier Transform (FFT) of a sequence of measured arterial pulse; (b) an example of the raw arterial pulses taken under 1.0 psi (51.7mmHg) at CGC positions; (c) CGC pulse signals after removal of the baseline-drift of the signals shown in Figure 6(b).

First, we applied a fast Fourier transform (FFT) to the raw data and obtained the frequency spectral distribution, which shows the amplitude of pressure as a function of frequency. As shown in Figure 6(a), the FFT reveals that most of the signal energy lies within the low-frequency ranges, especially below 2 Hz. As mentioned, this low-frequency signal may be due to the volunteer’s physical movement and normal variation in the systolic and diastolic pressure. The other two peak frequencies are the systolic and diastolic peaks of the pulse. If we apply classical wave theory, the beat frequency of the pulse will be the same as the difference between these 2 peak frequencies. The signal without baseline wandering, obtained by the removal of low-frequency noise is shown in Figure 6(c).

## Results

### Trial Experiments

Professor Jiangang SHEN from University of Hong Kong, a TCM expert, conducted sphygmopalpation and performed pulse pattern diagnosis on 3 volunteers. Their pulses were recorded using our PRH and processed as mentioned above. These volunteers were confirmed to have no cardiac abnormalities in the past and showed different waveforms (i.e., they were diagnosed as HUA, XI and CHEN). These pulses can be classified as different constitutions and Yin-Yang statuses in TCM diagnostics although the individuals’ regular arterial pulse without any changes in their personal health from the perspective of Western medicine. In the perspective of TCM, the pulse conditions can have relatively variability to different individuals with their constitutions even they are considered to be healthy according to Western medicine examination. In our study, for each person, we collected 5 minutes of pulse data at each CGC position and FZC pressure. As the sampling rate of the tactile sensor is 50 Hz, there are 15,000 data points (50 Hz x 60 sec x 5 min) per CGC position and FZC pressure. Thus, we obtained a total of 135,000 (15,000 x 9) data points, approximately 3,375 pulses per subject (assuming an average of 40 data points per pulse, which depends on the period of the pulse generated from every subject). Figure 7 (a) - (c) shows the pulse data processed after removing baseline wandering and 3D color contour map transformation.

**Figure 7:**
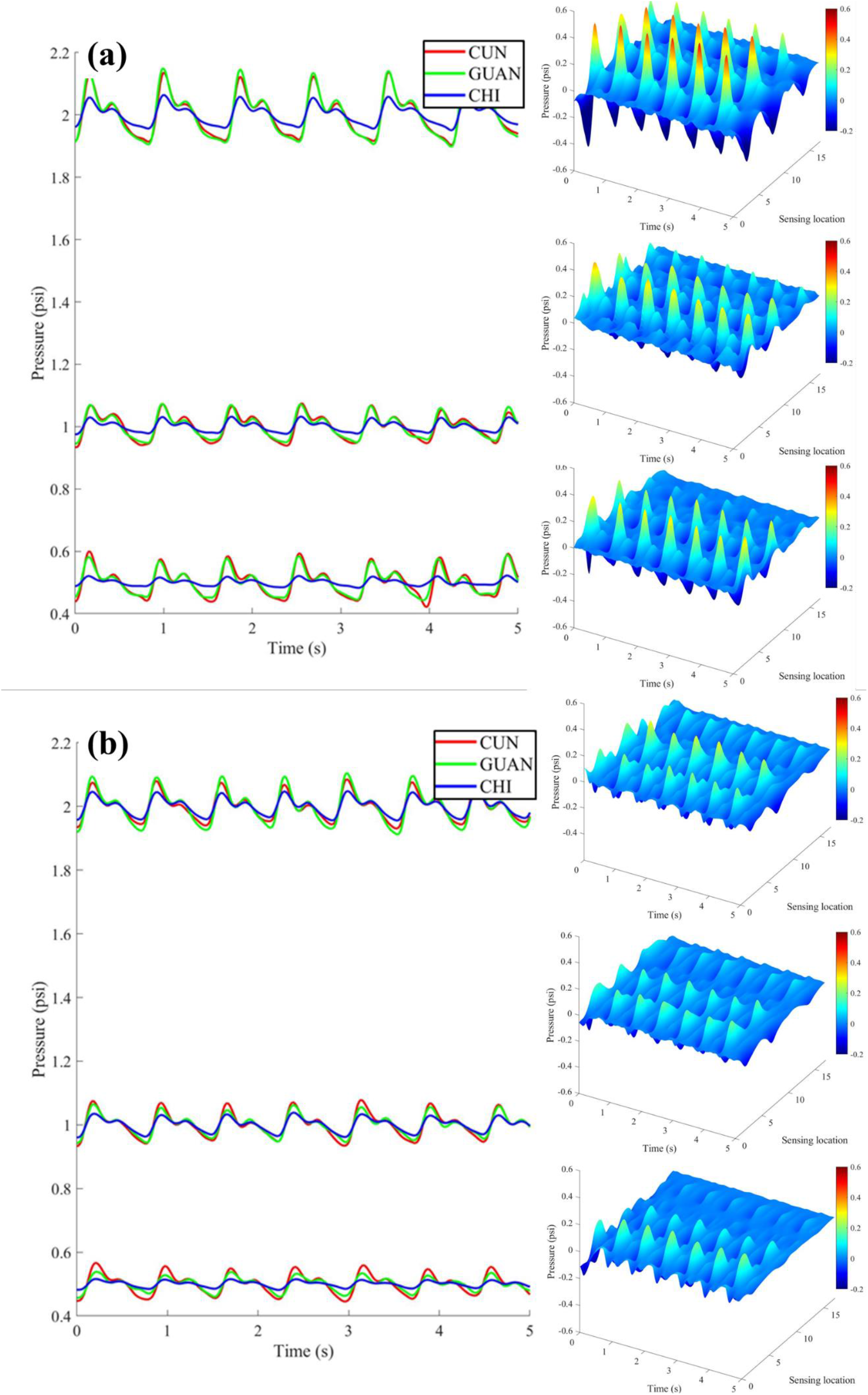
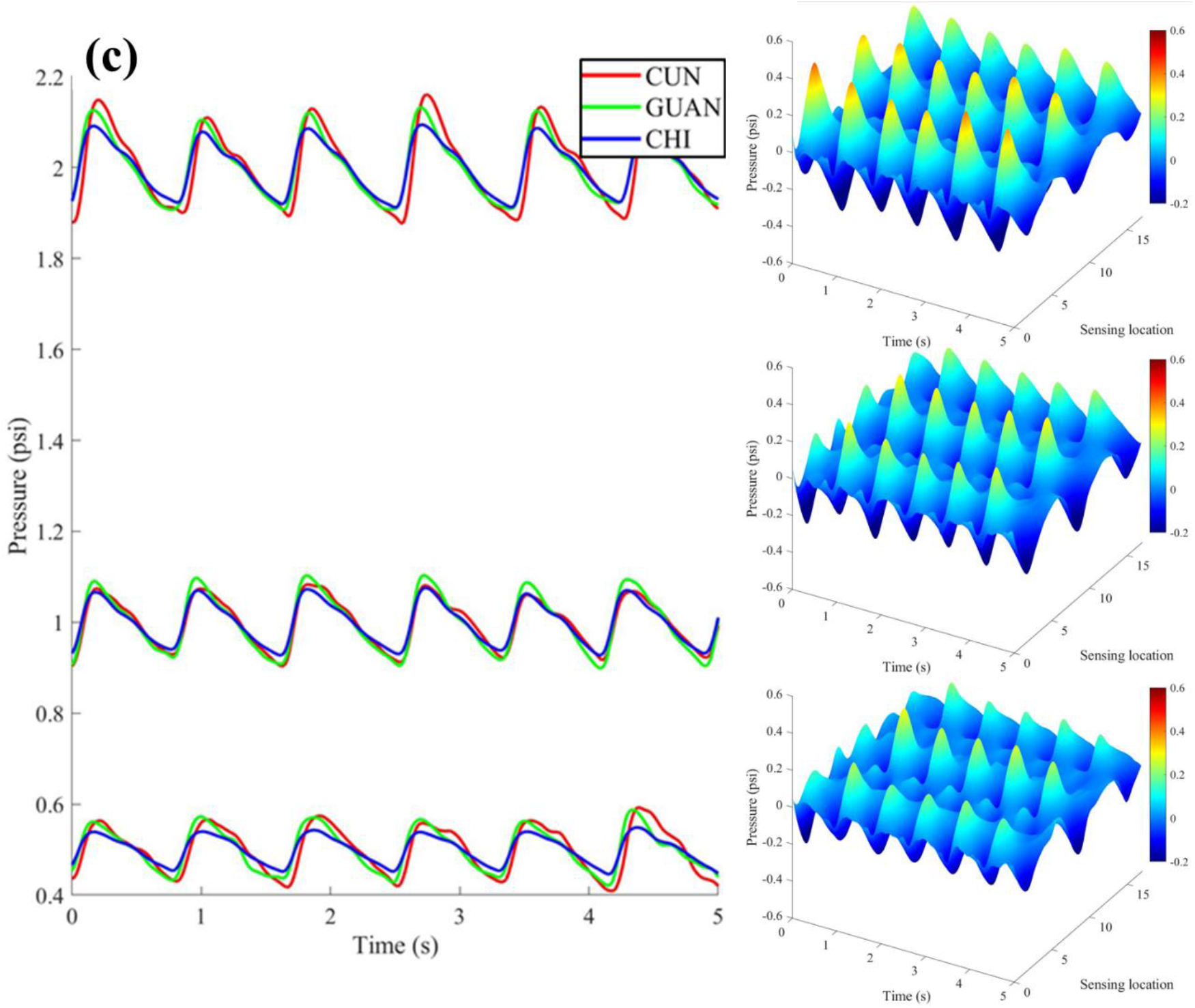
Pulse data of volunteers under 3 levels of applied pressure and 3 locations along with its 3D color contour map: (a) subject 1 diagnosed with HUA; (b) subject 2 diagnosed with XI; (c) subject 3 diagnosed with CHEN.

### Pulse Waveform Classification Using Deep Learning Algorithm

The collected 3D contour maps in Figure 3 consist most of the information including spatial, temporal and pressure information of one’s arterial pulse. To classify such a high-dimensional data, deep learning is the most suitable tool. Hence, convolutional neural network (CNN) was chosen as the analytic tool to classify pulse waveforms from the 3 volunteers. Before feeding the 3D contour maps into the CNN, we had to convert them into 2D images, namely “X-ray images” of human pulses. The detailed procedure of forming a grayscale 2D image is shown in Figure 8. The rows and columns of an 2D image, which is 54-by-54 pixels, represent the selected sensing location (i.e., sensing location 0 to 17 on the selected plane as shown in Figure 3) and time (i.e., 0.02 s (sampling frequency of our sensor) × 54 = 1.08s), respectively. These converted grayscale images were treated as inputs for the CNN, as shown in Figure 9, which has 3 outputs for classifying 3 different waveforms using the architecture listed in Table III. The pulses collected, as mentioned in last section, from the 3 volunteers can be converted into a total of 735 grayscale images. Each volunteer has 245 grayscale images which were divided randomly into three sets, namely, training, validation and testing, in the ratio of (0.6:0.3:0.1), respectively. The validation and testing accuracies of our CNN designed for arterial pulse pattern recognition were 98.7% and 84.2%, respectively, and its corresponding training progress is shown in Figure 10.

**Figure 8:**
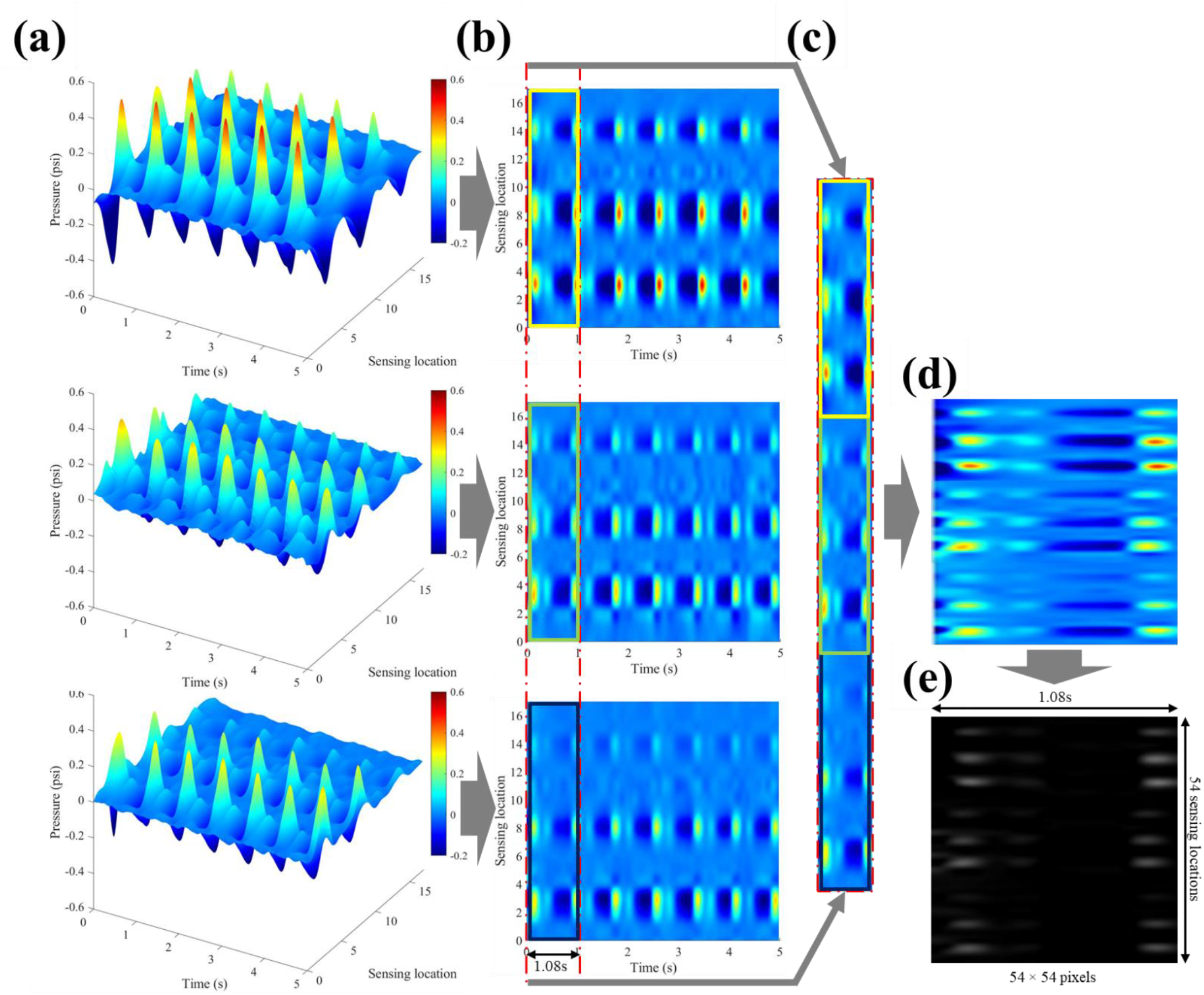
(a) 3D colour contour maps of an arterial pulse taken at CGZ location under FZC applied pressure by using the process described in Figure 3; (b) 2D projection of the time-location plane followed by image segmentations of the time-axis to crop a 18pixels-height (axis of sensing location) and 1.08s-width image sequentially for 3D colour contour maps in Figure 9(a); (c) combining the cropped images in Figure 10(b) together to form a 54-by-54 pixels image (height: 18pixels × 3 applied pressures = 54pixels; width: 1.08s × 50Hz =54pixels); (d) image reshaping and; (e) grayscale transfer to form an input image for the CNN.

**Figure 9:**
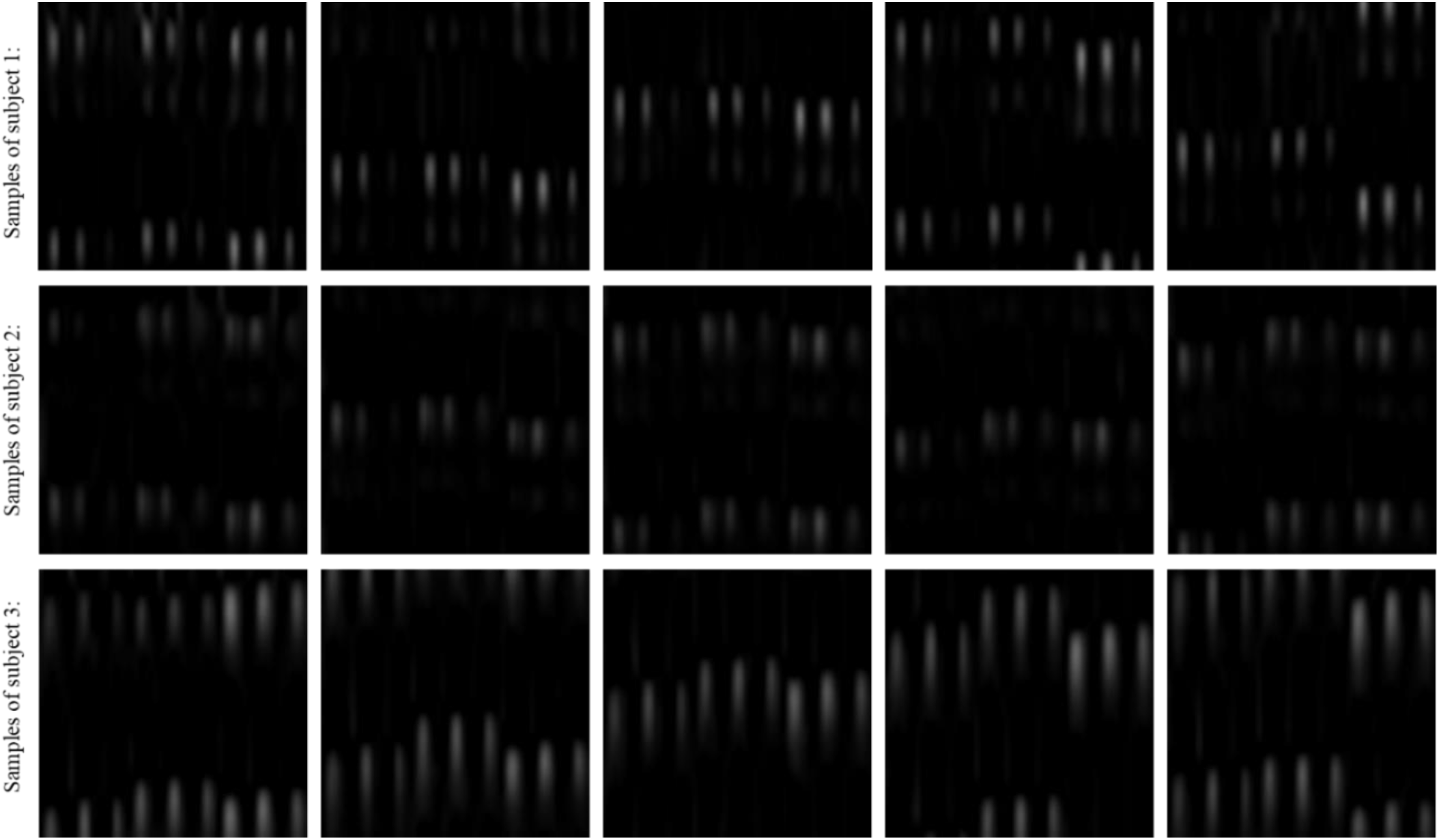
Input samples of the 3 subjects.

**Figure 10:**
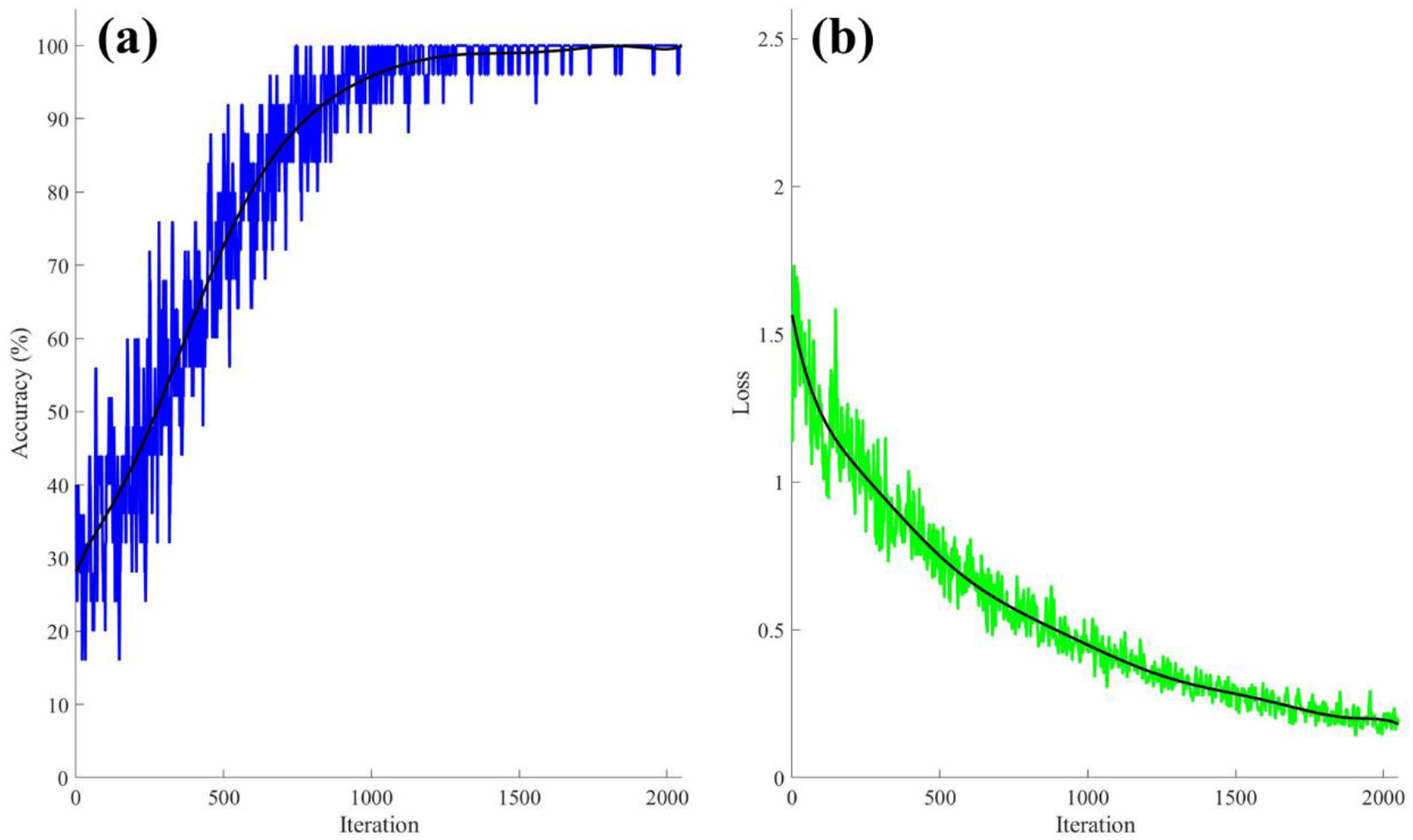
Training progress of the CNN: (a) classification accuracy on each individual mini-batch; (b) cross entropy loss of the training progress.

**Table III:** CNN architecture summarized parameters.

| Table III: CNN architecture summarized parameters. |  |  |  |  |  |  |
| --- | --- | --- | --- | --- | --- | --- |
| Layer |  | Feature map | Size | Kernel size | Stride | Activation |
| Input | Image | 1 | 54×54 | - | - | - |
| 1 | Convolution | 6 | 50×50 | 5×5 | 1 | tanh |
| 2 | Average polling | 6 | 25×25 | 2×2 | 2 | tanh |
| 3 | Convolution | 16 | 20×20 | 6×6 | 1 | tanh |
| 4 | Average pooling | 16 | 10×10 | 2×2 | 2 | tanh |
| 5 | Convolution | 120 | 1×1 | 5×5 | 1 | tanh |
| 6 | Full connection | - | 84 | - | - | tanh |
| 7 | Full connection | - | 10 | - | - | tanh |
| Output | Full connection | - | 3 | - | - | SoftMax |

### Conclusion

We have demonstrated an arterial PSP for pulse classification via the TCM approach. It consists of a robot with 3 sensing fingers, which mimics a CMP performing palpation, for arterial pulse measurement and an CNN for pulse classification. TCM practitioners rely on 3 fingers and 3 applied pressures for palpation to enrich the acquisition of pulse information which greatly increase the data structure and complexity. Hence, CNN is superior to other machine learning models in patterns classification because of its capabilities to tackle with complex data structure. In order to quantify human pulse in diagnostic applications, we proposed a methodology of obtaining “X-ray” image of pulse information constructed based on the sensing data from 3 locations (CGC) and 3 applied pressures (FZC), which contains all arterial pulse information in both spatial and temporal spans, and which could be used as an input to a deep learning algorithm. Compared to conventional approaches that use 2D information of pulses collected from a signal point, the proposed “X-ray” images preserve much more pulse information. Our preliminary results show that this platform can classify 3 pulse wave patterns with validation accuracy of 98.7% in training and prediction accuracy of 84.2% in testing.

With further improvement of the platform as well as the algorithm, we aim to digitalize and classify the basic 28 pulse characteristics described in TCM. This classification will help to minimize differences in TCM pulse diagnosis due to human variance by providing scientific-based pulse data information. Our long-term goal is to develop a data-based transmission of TCM knowledge, which was previously conveyed only by word of mouth or written notes.

## Acknowledge

This project is funded by the Research Grants Council (JLFS - RGC-Joint Laboratory Funding Scheme: Project no. JLFS/E-104/18) and by the Health and Medical Research Fund (HMRF - Health and Health Services (formerly HHSRF): Project no. 17181811).

